# White matter tract-oriented deformation is dependent on real-time axonal fiber orientation

**DOI:** 10.1101/2020.09.01.271502

**Authors:** Zhou Zhou, August G. Domel, Xiaogai Li, Gerald Grant, Svein Kleiven, David Camarillo, Michael Zeineh

## Abstract

Traumatic axonal injury (TAI) is a critical public health issue with its pathogenesis remaining largely elusive. Finite element (FE) head models are promising tools to bridge the gap between mechanical insult, localized brain response, and resultant injury. In particular, the FE-derived deformation along the direction of white matter (WM) tracts (i.e., tract-oriented strain) has been shown to be an appropriate predictor for TAI. However, the evolution of fiber orientation in time during the impact and its potential influence on the tract-oriented strain remains unknown. To address this question, the present study leveraged an embedded element approach to track real-time fiber orientation during impacts. A new scheme to calculate the tract-oriented strain was proposed by projecting the strain tensors from pre-computed simulations along the temporal fiber direction instead of its static counterpart directly obtained from diffuse tensor imaging. The results revealed that incorporating the real-time fiber orientation not only altered the direction but also amplified the magnitude of the tract-oriented strain, resulting in a generally more extended distribution and a larger volume ratio of WM exposed to high deformation along fiber tracts. These effects were exacerbated with the impact severities characterized by the acceleration magnitudes. Results of this study provide insights into how best to incorporate fiber orientation in head injury models and derive the WM tract-oriented deformation from computational simulations, which is important for furthering our understanding of the underlying mechanisms of TAI.

## Introduction

Traumatic brain injury (TBI) is a critical public health problem associated with high mortality and morbidity, and enormous diagnosis and treatment costs. In the United States, the Center for Disease Control and Prevention reported that the annual number of TBI-induced emergency department visits was around 1.37 million, resulting in an estimated expense of 60.43 billion dollars.^1^ A meta-analysis of injury data from 23 European countries revealed an overall TBI incidence rate of about 235 per 100000 and an average fatality rate of about 11 per 100.^2^ In Europe in 2010, a cost of 33 billion euros associated with TBI was reported.^3^ Although global effort has been made to reduce the occurrence and alleviate the detriment of TBI, neither TBI-related socioeconomic burden nor improved treatment outcome of TBI patients has been achieved.^4, 5^

This TBI-related dilemma may be partially due to the clinical difficulties of TBI assessment and diagnosis, which may be particularly true for traumatic axonal injury (TAI).^6, 7^ TAI is a frequent form of TBI.^8^ The clinical outcome of axonal injury exhibits a broad spectrum of cognitive impairment and pathological severity, ranging from mild levels, that can be associated with a transient alteration in consciousness and subtle neurological complaints, to severe forms, that can lead to prolonged coma accompanied by damage throughout the white matter (WM) of the brain.^9, 10^ Clinical detection of axonal injury, especially mild axonal injury, is difficult because the brain often exhibits no identifiable structural alteration based on conventional neuroimaging techniques (e.g., computed tomography and magnetic resonance imaging).^11–13^

In parallel, *in vivo, ex vivo*, and *in vitro* experimental models have been used to understand the mechanisms of axonal injury and define appropriate criteria and corresponding thresholds for TAI prediction by relating controlled mechanical inputs to resultant morphological damage or functional alteration of experimental tissue.^4, 14–16^ One of the most anatomically relevant mechanisms for TAI seems to be the tensile elongation of axonal fibers,^17–23^ which instigates immediate or delayed disconnections between neurons within WM tracts and subsequent large-scale neurological deficits. Based on this mechanism, any criteria related to the deformation of WM axonal fibers may be candidates for TAI prediction.

As an alternative to experiments, finite element (FE) head model is a numerical surrogate of the human head that can emulate the geometrical profiles and mechanical behavior of various intracranial components, as well as the interface condition among them. In recent decades, FE head models have been increasingly used to fill the knowledge gap between external loading, localized brain response, and resultant injury.^24–29^ Partially inspired by the aforementioned mechanism that axonal elongation triggers TAI occurrence, various endeavors have been made recently to simulate deformation along WM fiber tracts (i.e., the tract-oriented strain).^30–44^ Of note, the tract-oriented strain is synonymous with axonal strain or fiber strain, which are both terms used in multiple studies.^39, 45^ As was common in these efforts, an advanced neuroimaging technique called diffusion tensor imaging (DTI)^46^ was exploited to delineate orientation information of fiber tracts and this orientation information can be combined with FE models. Three different techniques have been proposed thus far to integrate the orientation of WM tracts from DTI into TBI analysis: (1) Chatelin and colleagues^47^ developed a post-processing technique to obtain the tract-oriented strain by projecting the strain tensors from pre-computed simulations onto the direction of fiber tracts; (2) Cloots and colleagues^32^ and Giordano and coworkers^36^ instead assigned a hyper-viscoelastic fiber-reinforced anisotropic material to the brain with the axonal fiber orientation integrated into the constitutive law to inform the fiber reinforcement; (3) Garimella and Kraft^35^ proposed another numerical technique to model whole-brain fiber tractography explicitly using an embedded element method. Despite these aforementioned technical disparities along with other model-specific computational choices, it has been repeatedly reported by several independent groups that tract-oriented strain was a more appropriate predictor for TAI than its isotropic counterparts (e.g., maximum principal strain, cumulative strain damage measures (CSDM)).^37, 39, 41, 45, 48, 49^

Although tract-oriented strain is a spatiotemporal-dependent variable and highly sensitive to the temporal variation of axonal fiber orientation,^50, 51^ the evolution of fiber orientation in time during an impact, as well as its potential influence on the tract-oriented strain, has rarely been reported in the literature. Only a few studies^38, 41^ clarified that both the strain tensors from the pre-computed simulations and fiber orientation from DTI were commonly transformed into a skull-fixed coordinate system and then the strain tensors were projected onto the time-invariant fiber direction to obtain the tract-oriented strain. Such a transforming operation accounted for the variation of fiber orientation in relation to skull movement within the global coordinate system. Nevertheless, the axonal fiber may still be reoriented within the skull-fixed coordinate system owing to the brain deformation and brain-skull relative motion during impact.^41, 48^ Thus, real-time tracking of the fiber direction and employing temporal fiber orientation to inform the calculation of the tract-oriented strain have yet to be properly considered.

The aim of this study is to investigate real-time fiber orientation during head impacts and its resultant influence in the prediction of WM tract-oriented deformation. To achieve this, an embedded element approach was leveraged to monitor the temporal direction of fiber tracts in a previously established three-dimensional (3D) head model, from which the deviation of fiber orientation during impacts was quantified. By comparing two types of tract-oriented strains obtained via the projection of strain tensors in pre-computed simulations onto the temporal fiber orientation and its static counterpart from DTI, the influence of real-time fiber orientation on the tract-oriented strain was evaluated. We thus tested the hypothesis that the real-time fiber orientation affects both the direction and magnitude of the tract-oriented strain, and consequently the distribution and extent of WM with high deformation along the fiber tracts.

## Methods

### Finite element head model

The FE model, referred to as the Baseline-model, used in this study was previously developed at the Royal Institute of Technology (KTH) in Stockholm by Kleiven^52^ using LS-DYNA. The Baseline-model includes the scalp, skull, brain, meninges (i.e., dura mater and pia mater), intracranial membranes (i.e., falx and tentorium), cerebrospinal fluid (CSF), eleven pairs of the largest parasagittal bridging veins, and a simplified neck with the extension of the spinal cord (Figure 1). The brain elements were further grouped to represent the primary brain components, including the cerebral gray matter (GM), cerebral WM, corpus callosum, thalamus, brainstem, midbrain, cerebellar GM, cerebellar WM, and ventricles. Detailed information regarding the geometry discretization and material choice for each head component in the Baseline-model are available in previous studies by Kleiven.^52, 53^ It is worth mentioning that the brain was simulated as an isotropic medium in the current study owing to the lack of consensus regarding the mechanical anisotropy of the brain.^54^ Specifically, a second-order hyperelastic constitutive law was used to describe the nonlinear behavior of the brain tissue with additional linear viscoelastic terms to account for its rate dependence. Responses of the Baseline-model have shown good correlation with experiments of brain-skull relative motion,^25, 55^ intracranial pressure,^56^ skull fracture,^57^ and brain strain.^58^

**Figure 1.**
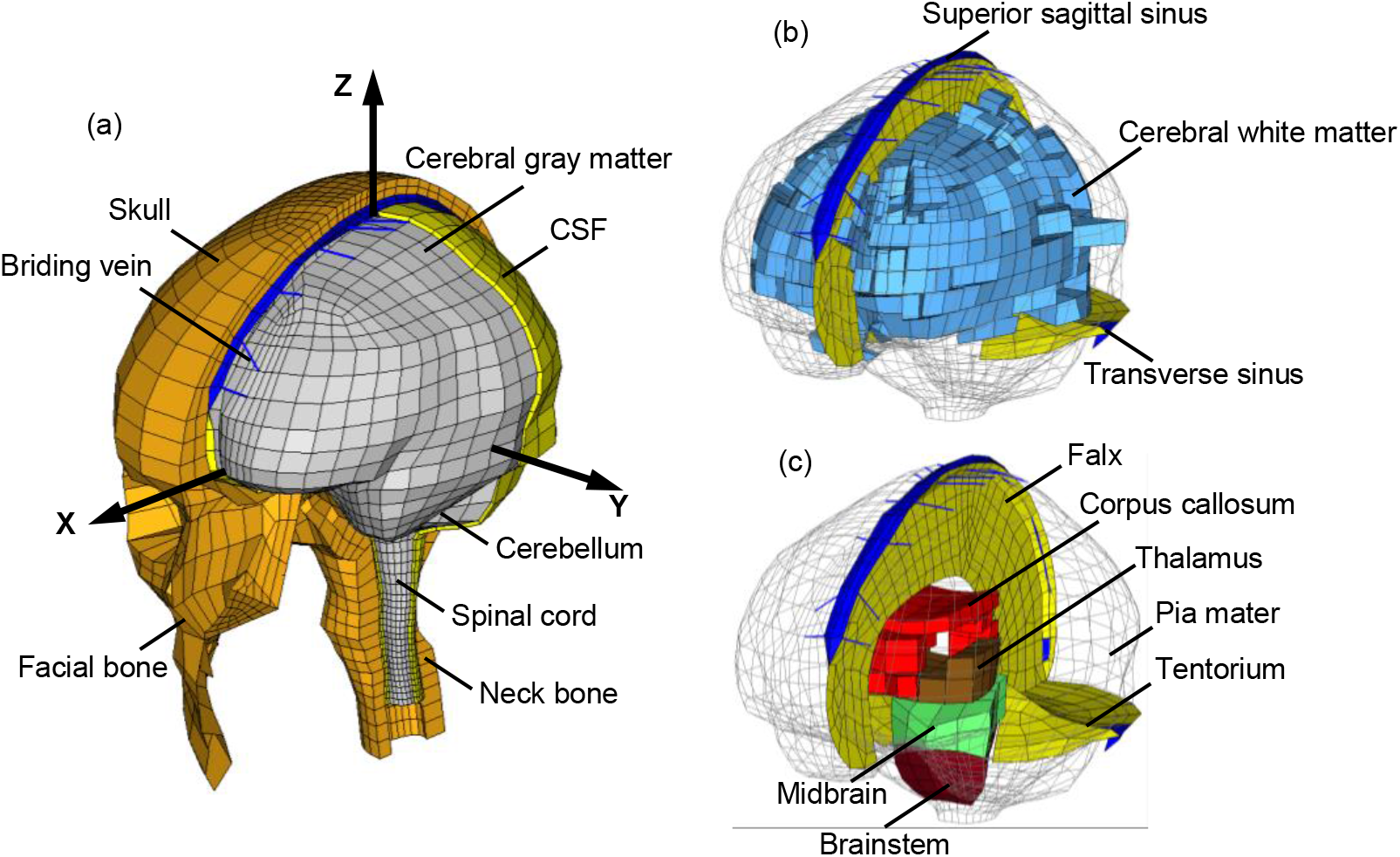
(a) Isometric view of the finite element model of the human head with the skull open to expose the CSF and brain. A skull-fixed coordinate system and corresponding axes are illustrated with the origin at head’s center of gravity. (b) Isometric view of cerebral white matter and intracranial sinuses. (c) Isometric view of deep brain structures and membranes.

### Computation of the tract-oriented strain with static fiber orientation

To obtain the deformation of brain tissue along the primary direction of fiber tracts, a post-processing technique was previously implemented in the Baseline-model by coupling the element-level strain response with fiber orientation information derived from DTI.^36, 38^ The averaged tractography was obtained from the ICBM DTI-81 atlas,^59^ which was aligned with the brain mask of the Baseline-model via an affine registration. For a given WM element in the Baseline-model, corresponding voxels in the DTI that occupied the same space as that of the WM element were identified. Diffusion information of these identified voxels were then averaged using equation (1), in order to more heavily weight the voxels closer to the WM element centroid:

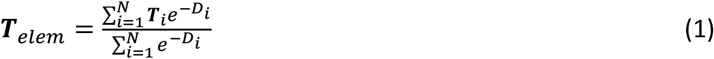

where ***T**_elem_* is the mean diffusion tensor calculated for each WM element; *N* is the number of selected DTI voxels for a given WM element; ***T**_i_* and *D_i_* are the diffusion tensor in the DTI voxel, and the normalized distance from the center of the DTI voxel to the centroid of the WM element, respectively.

The principal eigenvector (***v**_o_*) of ***T**_elem_* was obtained for each WM element and are hereafter referred to as the original fiber orientation. Note that ***v**_o_* was expressed with reference to the skull-fixed coordinate system (i.e., the coordinate system in Figure 1 (a)). The strain tensor for each WM element at each timestep was derived from the LS-DYNA solver and then transformed into the skull-fixed coordinate system, similar to the approach adopted by Sullivan and colleagues.^41^ The tract-oriented strain was computed according to equation (2):

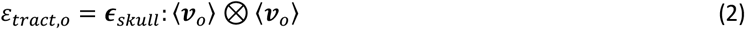

where *ε_tract,o_* is the tract-oriented strain projected along ***v**_o_*; *ϵ_skull_* is the Green-Lagrange strain tensor of WM elements in the skull-fixed coordinate system.

Details regarding the coupling protocol between DTI and FE brain model and computation of *ε_tract,o_* are available in previous studies by Giordano and coworkers.^36, 38^ Given that ***v**_o_* remained static during the whole post-processing procedure, the potential deviation of fiber orientation induced by brain-skull relative motion and brain deformation during the impact was not considered.

### Implementation of embedded element method for real-time fiber orientation

Given the importance of the fiber orientation on the post-processing technique, an embedded element method was implemented in the Baseline-model to monitor the real-time fiber orientation during impact. For each WM element (serving as a master element), the inserted DTI-derived fiber orientation was concretely represented by an embedded truss element, which oriented in the same direction as that of ***v**_o_*. As illustrated in Figure 2 (c), one node of the truss element was located at the centroid of its master element and the other fell within the boundary of the master element. Note that the truss element only served as an auxiliary to monitor the real-time fiber orientation and no extra stiffness and other physical properties were added by the truss element. Thus, the material properties of the truss element were represented by a null constitutive model in LS-DYNA with nominal density and cross-sectional area (Table 1).^60^ Using this modeling strategy, the temporal orientation of the truss element and strain response of its master element were independent of the length of the embedded truss element, as detailed in Appendix A. Nodal motion of a given truss element was exclusively governed by its master element based on equation (3) and (4):^61^

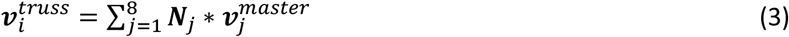

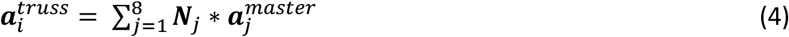

where 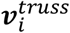 and 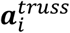 are the velocity vectors and acceleration vectors of the nodes in the truss element, respectively; *i* = 1,2, which corresponds to the two nodes of a truss element; *N_j_* is the shape function of the master element that hosts the truss element; 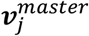 and 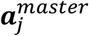 are the velocity vectors and acceleration vectors of the 8 nodes of the hexahedral elements (i.e., master element), respectively.

**Figure 2.**
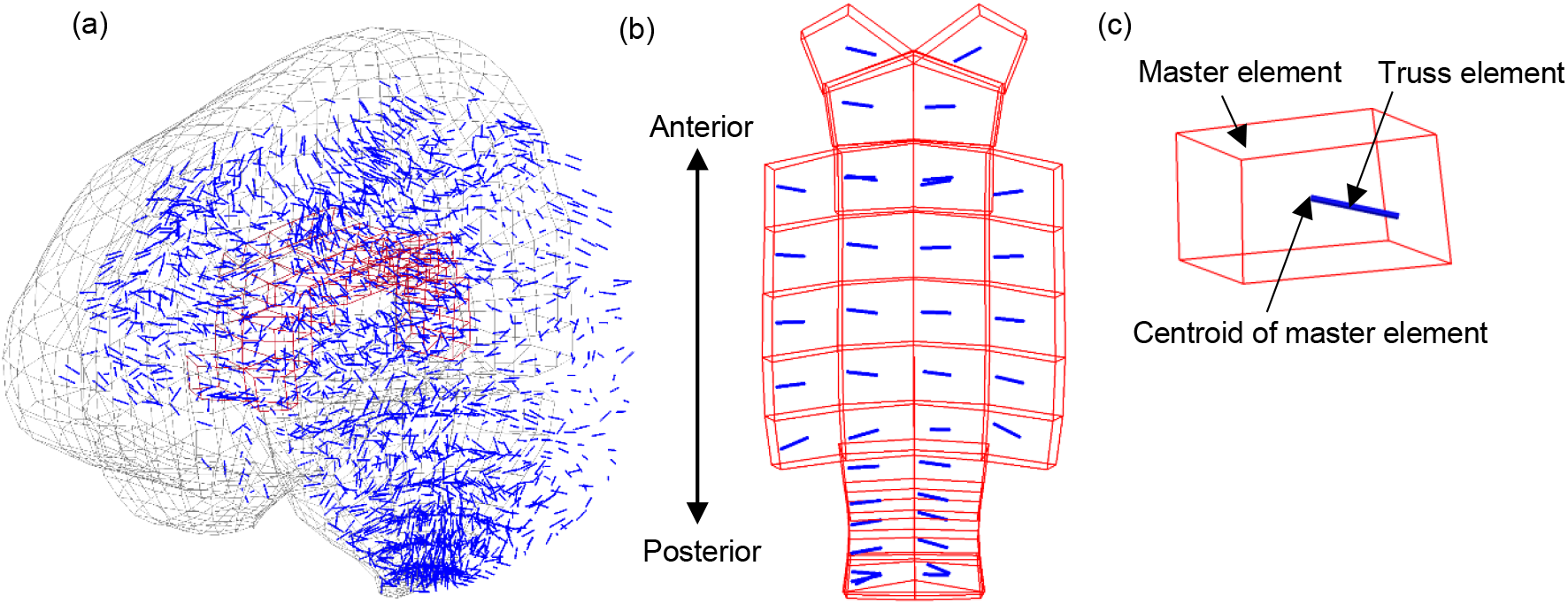
(a) Isometric view of the brain (in gray) with embedded truss elements (in blue). For better illustration, half of the brain is masked with the corpus callosum highlighted in red. (b) Top view of the truss elements within the corpus callosum. (c) Isometric view of a representative element in the corpus callosum with a truss element embedded.

**Table 1.** Material properties for the embedded truss elements.

| Element type | Cross-sectional area (m<sup>2</sup>) | Constitutive model | Density (kg/m<sup>3</sup>) | Reference |
| --- | --- | --- | --- | --- |
| Truss | 1 e-11 | MAT_NULL | 1 e-11 | <sup>60</sup> |

These two equations constrained both acceleration and velocity of the truss element to its master element in all directions. Consequently, potential variation in fiber orientation associated with brain deformation and brain-skull relative motion during the impacts were reflected by the temporal direction of the truss element, which was updated at each solution cycle of the time-marching simulation. Once the simulation was completed, both the strain tensors for all WM elements and the orientation of the embedded truss elements were obtained for each timestep and then transformed into the skull-fixed coordinate system. Instead of equation (2), equation (5) was used to calculate the updated tract-oriented strain as follows:

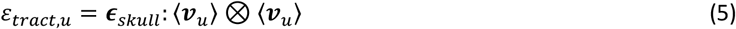

where *ε_tract,u_* is the tract-oriented strain projected along ***v**_u_*. Here, ***v**_u_* is the updated truss orientation output from LS-DYNA per each timestep and is expressed with reference to the skull-fixed coordinated system. Under the undeformed configuration, ***v**_u_* equaled to ***v**_o_* for all WM elements. This modified model is referred to as Truss-embedded-model hereafter.

### Loading conditions

At Stanford University, instrumented mouthguards have been developed to measure six-degree-of-freedom head kinematics during in-game head impacts to athletes.^62, 63^ Using these instrumented mouthguards, over 500 head impacts in football have been video confirmed.^64^ In the current study, two concussive impacts and ten randomly selected subconcussive impacts^65^ were imposed to both the Baseline-model and Truss-embedded-model, respectively, to unravel the deviation of fiber orientation in realistic impacts and its influence on the computation of the tract-oriented strain. Of these two concussive impacts, one resulted in the athlete suffering loss of consciousness, while the other was self-reported with post-concussive symptoms.^64^ A summary of the acceleration peaks of the 12 selected realistic impacts is listed in Table 2 with detailed acceleration profiles of the two concussive impacts exemplified in Figure 3.

**Table 2.** Peaks of translational acceleration, rotational acceleration, and injury severity of the 12 cases considered in this study

| Case ID | Peak translation acceleration (g) |  |  |  | Peak rotational acceleration (krad/s<sup>2</sup>) |  |  |  | Injury severity |
| --- | --- | --- | --- | --- | --- | --- | --- | --- | --- |
|  | X | Y | Z | Magnitude | X | Y | Z | Magnitude |  |
| Case 1 | -40.6 | 100.4 | -63.4 | 106.1 | 12.89 | -3.06 | -3.24 | 12.95 | Concussion |
| Case 2 | -61.1 | -57.8 | -45.8 | 84.2 | 4.21 | 5.14 | -1.84 | 6.19 | Concussion |
| Case 3 | -10.3 | 11.6 | -7.2 | 15.6 | 0.66 | -0.81 | -1.86 | 1.98 | Sub-concussion |
| Case 4 | -44.7 | 32.2 | 46.1 | 57.5 | 1.97 | 3.88 | 1.81 | 4.42 | Sub-concussion |
| Case 5 | -34.0 | 18.7 | -22.5 | 42.0 | 0.64 | -1.61 | -0.55 | 1.65 | Sub-concussion |
| Case 6 | -14.3 | 10.7 | 15.5 | 17.6 | 0.61 | 1.87 | -0.95 | 2.05 | Sub-concussion |
| Case 7 | -52.9 | -22.6 | -36.5 | 63.9 | 1.69 | -5.22 | 1.51 | 5.27 | Sub-concussion |
| Case 8 | -25.4 | 5.2 | -15.0 | 28.9 | -0.42 | -1.66 | 0.62 | 1.70 | Sub-concussion |
| Case 9 | 33.5 | -13.3 | 28.8 | 44.8 | 0.68 | 4.42 | -0.32 | 4.45 | Sub-concussion |
| Case 10 | -46.0 | 5.9 | 30.5 | 46.3 | 0.57 | -2.52 | 0.53 | 2.58 | Sub-concussion |
| Case 11 | -38.3 | -15.4 | -39.6 | 55.6 | 1.02 | -5.21 | 0.70 | 5.22 | Sub-concussion |
| Case 12 | 23.3 | 14.0 | 25.3 | 32.8 | 1.29 | 2.83 | -0.82 | 2.91 | Sub-concussion |

**Figure 3.**
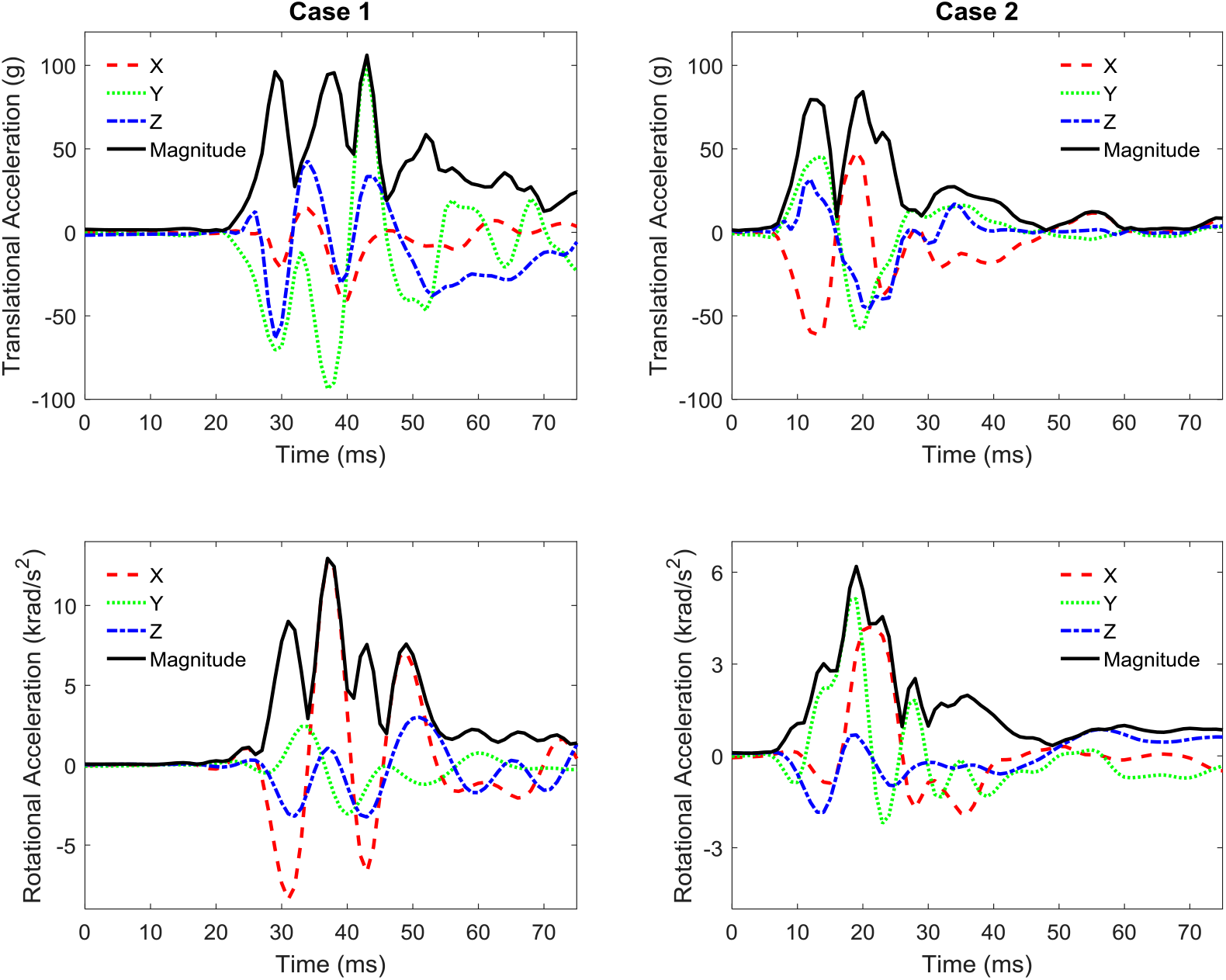
Head model loading condition for the two concussive cases. Note that the X, Y, and Z axes are the same as those in the skull-fixed coordinate system in Figure 1 (a).

To test the potential dependency of fiber orientation variation and tract-oriented strain on impact severity, 4 artificial loadings were generated by halving and doubling each component of acceleration profiles of Case 1 and Case 2 (Figure 3). These 4 artificial loadings were imposed to both the Baseline-model and Truss-embedded-model.

In all the simulations, translational and rotational accelerations were applied to a node located at the head’s center of gravity and constrained to a rigidly modelled skull. A selectively reduced integration scheme was used for all the brain components and CSF. A typical simulation of 90 ms required about 10 hours for the Baseline-model and 11 hours for the Truss-embedded-model on a local Linux cluster using a single processor. The model responses were output at every 0.1 ms.

### Post-processing and data analysis

To quantify the variation of fiber orientation during the impacts, the relative angle between the two tract-oriented strain measures (*θ_<ε_tract,o_,ε_tract,u_>_*) was calculated for each WM element at each timestep. To ascertain the effect of fiber orientation variation on the computation of the tract-oriented strain, *ε_tract,o_* and *ε_tract,u_* were calculated for the Baseline-model with the fiber orientation originally obtained from DTI (i.e., equation (2)) and the Truss-embedded-model with the fiber orientation updated at each time step (i.e., equation (5)), respectively. The peak values of *ε_tract,o_* and *ε_tract,u_* were denoted by 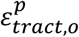 and 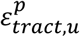, respectively. To exclude the potential numerical the 95th percentile 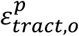 and 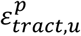 (referred to as 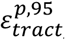 and 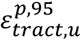, respectively) were presented for all 12 impacts, similar to the strategies in previous studies.^66, 67^ To statistically determine the effect of the real-time fiber orientation on the tract-oriented strain, a Wilcoxon matched-pairs signed-rank test was performed to examine the difference between 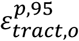 and 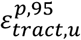. The threshold for significance was p<0.05.

To unravel the effect of impact severity (characterized by acceleration magnitude) on fiber orientation variation and tract-oriented strain, responses of the two realistic concussive impacts were compared to their counterparts with artificial loadings that possessed the same shapes of acceleration profiles but varied in magnitudes as those of two realistic concussive impacts. A detailed summary of the FE-derived variables is listed in Table 3.

**Table 3.** Summary of the FE-derived variables.

|  |  |
| --- | --- |
| $\varepsilon_{tract,o}$ | Tract-oriented Green-Lagrange strain with fiber orientation originally obtained from DTI |
| $\varepsilon_{tract,u}$ | Tract-oriented Green-Lagrange strain with fiber orientation updated at each time step |
| $\theta_{<\varepsilon_{tract,o}, \varepsilon_{tract,u}>}$ | Absolute value of the angle between the two tract-oriented strain measures |
| $\varepsilon_p$ | First principal Green-Lagrange strain |
| $\varepsilon_{tract,o}^p$ | The accumulated peak $\varepsilon_{tract,o}$, regardless of the time of occurrence |
| $\varepsilon_{tract,u}^p$ | The accumulated peak $\varepsilon_{tract,u}$, regardless of the time of occurrence |
| $\varepsilon_p^p$ | The accumulate peak $\varepsilon_p$, regardless of the time of occurrence |
| $\varepsilon_{tract,o}^{p,95}$ | 95<sup>th</sup> percentile of $\varepsilon_{tract,o}^p$ for the whole white matter elements |
| $\varepsilon_{tract,u}^{p,95}$ | 95<sup>th</sup> percentile of $\varepsilon_{tract,u}^p$ for the whole white matter elements |
| $\theta_{<\varepsilon_p, \varepsilon_{tract,o}>}$ | Absolute value of the angle between $\varepsilon_p$ and $\varepsilon_{tract,o}$ |
| $\theta_{<\varepsilon_p, \varepsilon_{tract,u}>}$ | Absolute value of the angle between $\varepsilon_p$ and $\varepsilon_{tract,u}$ |

## Results

To quantify the variation of fiber orientation during the impacts, the relative angle between the two tract-oriented strain measures was extracted. The time-history curves of maximum *θ_<ε_tract,o_,ε_tract,u_>_* in six brain subregions are plotted in Figure 4 for the two concussive impacts. For Case 1, the maximum *θ_<ε_tract,o_,ε_tract,u_>_* ranged from 15°in the brainstem, to 31°in cerebral WM. For Case 2, the maximum *θ_<ε_tract,o_,ε_tract,u_>_* varied from 9° in cerebellar WM, to 25° in cerebral WM. Figure 5 (a) illustrates both the original and updated (blue and red, respectively) configurations of the truss elements embedded within the corpus callosum for Case 1. The element with maximum *θ_<ε_tract,o_,ε_tract,u_>_* within the corpus callosum (29°) is shown in the enlarged view. A similar illustration was presented for Case 2 in Figure 5 (b) with the thalamus as the representative region, in which the maximum *θ_<ε_tract,o_,ε_tract,u_>_* was 16°. Element-wise *θ_<ε_tract,o_,ε_tract,u_>_* peaks for all WM elements in all 12 impacts are summarized in Figure 6. The data were presented in the form of mean value ± standard deviation with the values ranging from 3.0° ± 1.9° in Case 5 to 10.2° ± 5.2° in Case 1.

**Figure 4.**
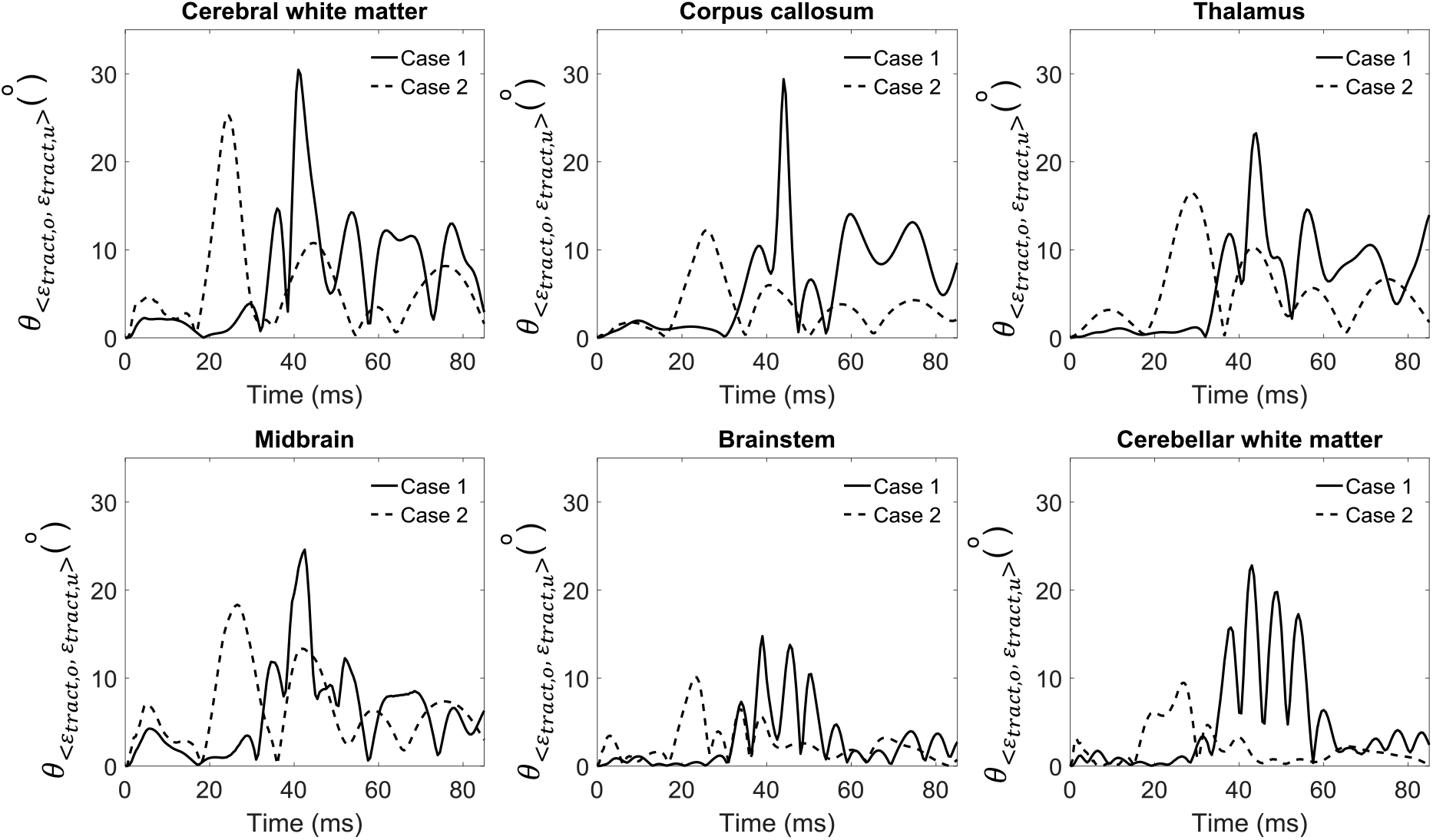
Time-history curves of maximum *θ_<ε_tract,o_,ε_tract,u_>_* in six brain subregions for two concussive impacts.

**Figure 5.**
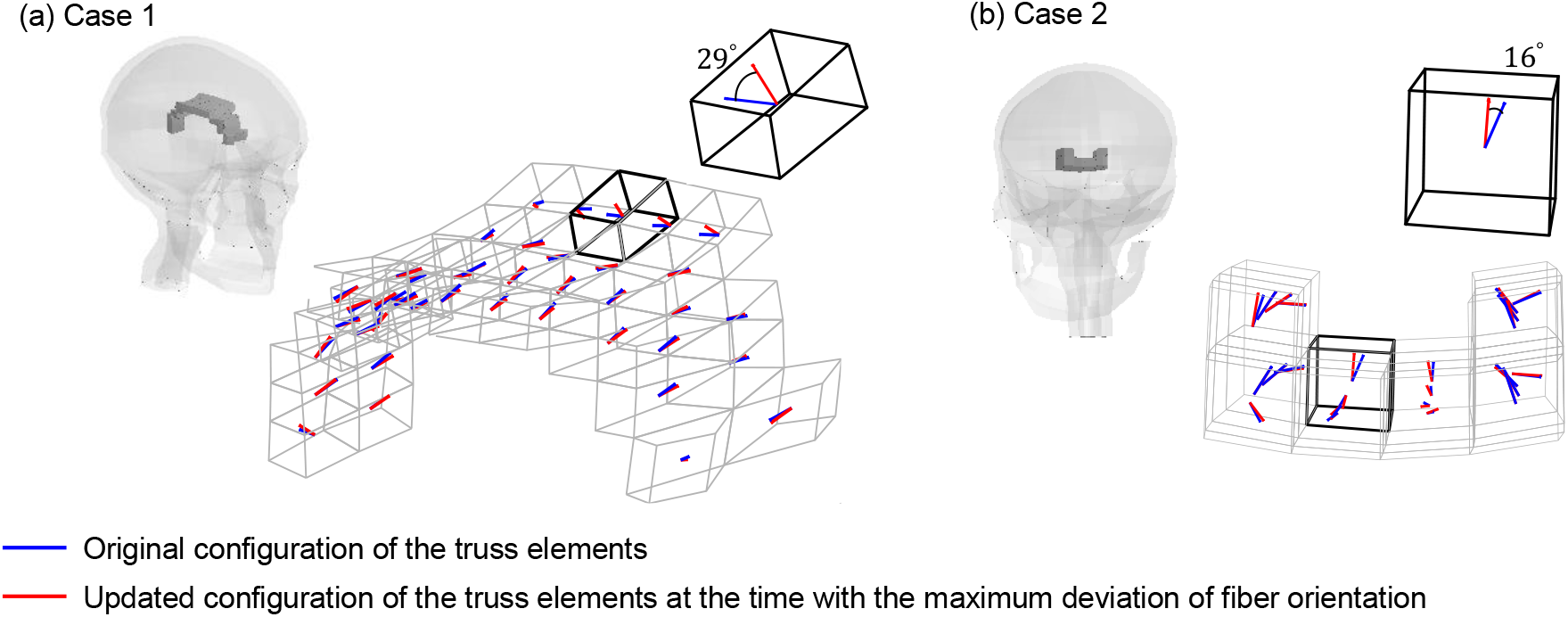
(a) Original and updated configurations of the truss elements embedded within the corpus callosum (in grey wire) for Case 1 with a whole-head thumbnail showing the view angle, and a zoomed view showing the element (in bold black wire) with the largest deviation of fiber orientation in the corpus callosum. (b) Original and updated configurations of the truss elements embedded within the thalamus (in grey wire) in Case 2 with a whole-head thumbnail showing the view angle, and a zoomed view showing the element (in bold black wire) with the largest deviation of fiber orientation. In both subfigures, the original configuration of the truss elements is in blue, while their updated counterparts in red. The master elements in their undeformed configurations are also illustrated in both subfigures.

**Figure 6.**
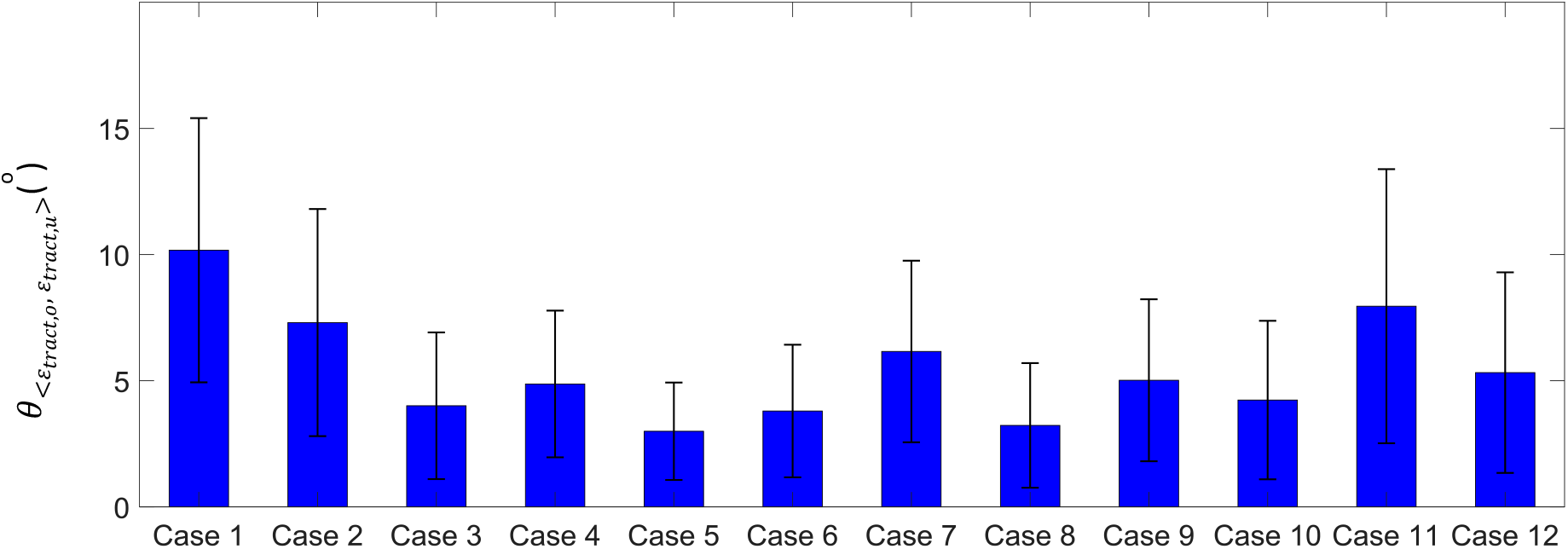
Summary of element-wise *θ_<ε_tract,o_,ε_tract,u_>_* peak for the 12 impacts in the form of mean value ± standard deviation.

To identify the effect of fiber orientation variation on the computation of the tract-oriented strain, both *ε_tract,o_* and *ε_tract,u_* were calculated for the 12 selected impacts. For the two concussive impacts, the differences between the 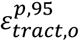 and 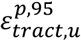 across all WM elements were 21.27% for Case 1 (Figure 7 (a)), and 18.60% for Case 2 (Figure 7 (c)). All WM elements with tract-oriented strain peaks over 0.14, a conservative threshold proposed by Bain and Meaney^68^ to induce morphological damage to WM, were additionally identified to calculate the volume ratio of injured WM to the whole WM in the FE model. By comparing the injured WM element based on the two types of tract-oriented strains, a generally more extended distribution of injured WM elements was noted for 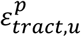 in both concussive cases (Figure 7 (b) and Figure 7 (d)). In Case 1, the volume ratios of injured WM were 18.91% for *ε_tract,o_* and 25.7% for 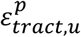. In Case 2, the volume ratios of injured WM were 3.42% to 6.12% for 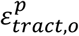 and 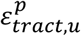, respectively. For the 10 subconcussive impacts, larger magnitudes were consistently noted for 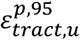 than 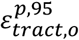 with the difference varying from 8.07% in Case 4 to 26.66% in Case 5 (Figure 7 (e)). A Wilcoxon matched-pairs signed-rank test performed between 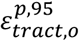 and 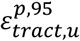 for all 12 impacts showed the difference between the two types of tract-oriented strains was significant (p=0.002).

**Figure 7.**
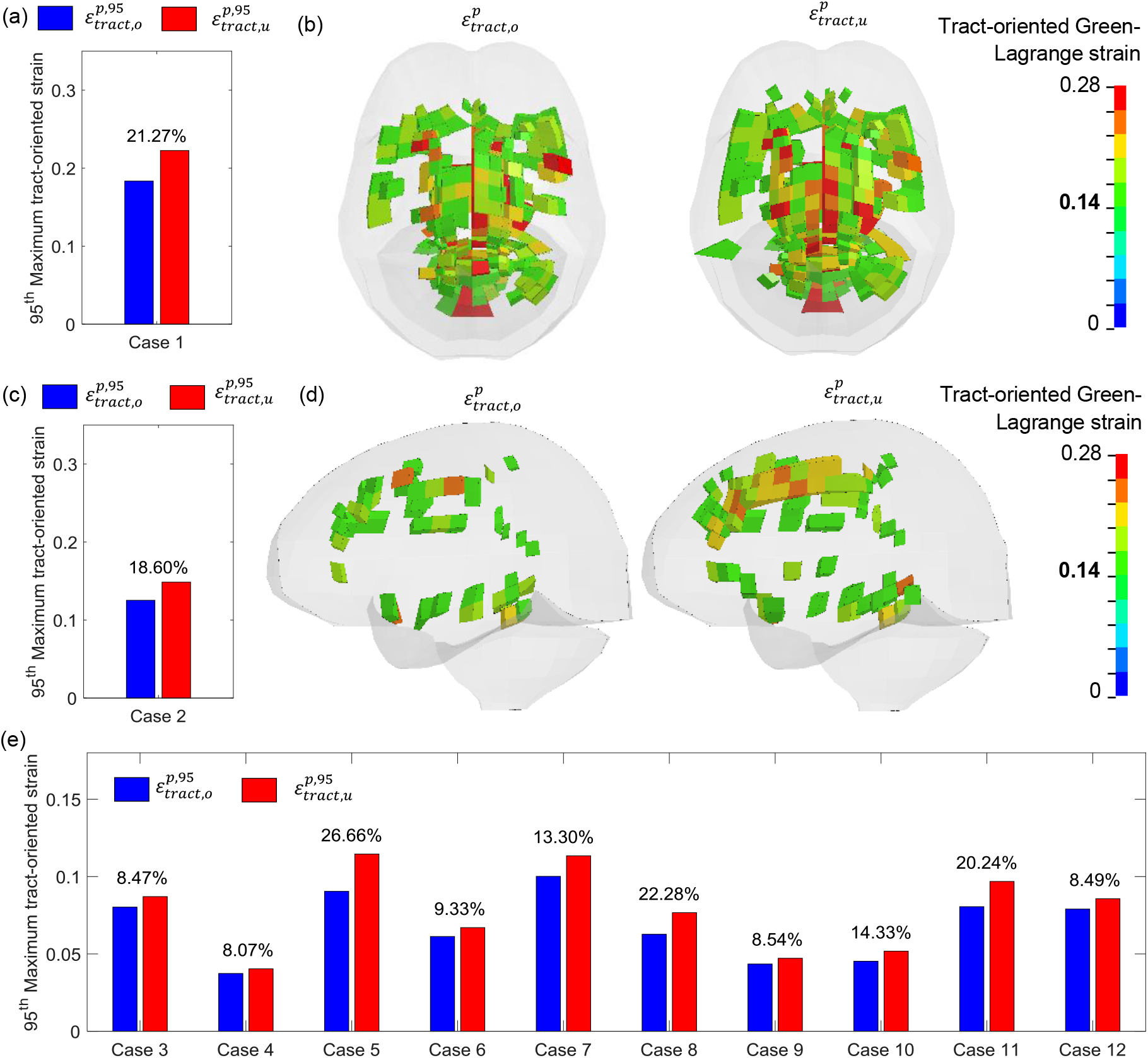
(a) Comparison of 95^th^ percentile maximum tract-oriented strain for Case 1. (b) Top view of white matter elements with tract-oriented strain peaks over 0.14 for Case 1. (c) Comparison of 95^th^ percentile maximum tract-oriented strain for Case 2. (d) Side view of white matter elements with tract-oriented strain peaks over 0.14 for Case 2. (e) Comparison of 95^th^ percentile maximum tract-oriented strain for 10 subconcussive cases. Note that Case 1 and Case 2 are concussive impacts. Percentage values for the difference between 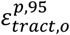 and 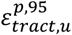 are listed in the bar figures.

To examine the relationship between the two types of tract-oriented strains and maximum principal strain, the time-history curves of these three strain measures, as well as their relative angles for one representative element derived from the two concussive impacts, were plotted (Figure 8). For Case 1, the representative element is the one with maximum fiber deviation in the corpus callosum (Figure 8 (a) and Figure 8 (c)). For Case 2, the representative element is the one with maximum fiber deviation in the midbrain (Figure 8 (b) and Figure 8 (d)). Regardless of impact cases, *ε_p_* was larger than the two types of tract-oriented strains, while *ε_tract,u_* was larger than *ε_tract,o_* (Figure 8 (a) and (b)). As plotted in Figure 8 (c) and (d), *θ_<ε_p_,ε_tract,u_>_* was generally smaller than *θ_<ε_p_,ε_tract,o_>_* for both impacts, which was particularly notable at the peaking time of *ε_tract,u_* as identified by the gray dash lines. It was indicated that the direction of *ε_tract,u_* was more aligned with that of *ε_p_*.

**Figure 8.**
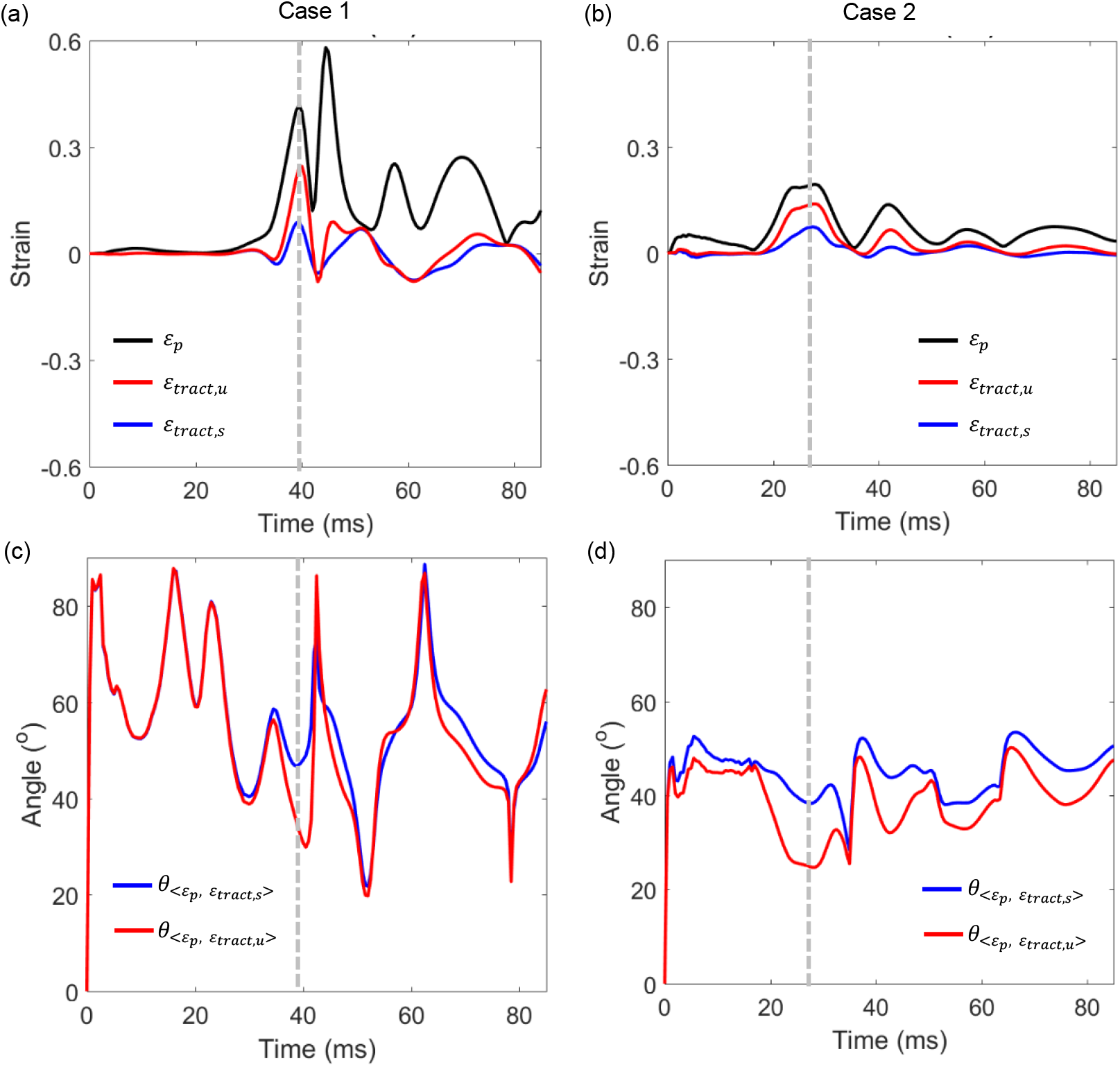
(a) Time history curves of *ε_p_, ε_tracto_*, and *ε_tract,u_* for one representative element in the corpus callosum with the maximum deviation of fiber orientation in Case 1. The relative angles between *ε_p_* and two tract-oriented strains plotted in subfigure (c). The grey dash lines in subfigure (a) and (c) identify the peaking time of *ε_tract,u_* in Case 1. (b) Time history curves of *ε_p_, ε_tract,o_*, and *ε_tract,u_* for one representative element in the midbrain with the maximum deviation of fiber orientation in Case 2. The relative angles between the *ε_p_* and the two tract-oriented strains plotted in subfigure (d). The grey dash lines in subfigure (b) and (d) identify the peaking time of *ε_tract,u_* in Case 2.

To evaluate the dependency of fiber orientation variation and tract-oriented strain on impact severity, responses of the two realistic concussive impacts were compared to their counterparts with artificial loadings that possessed the same shapes of acceleration profiles but varied in magnitudes as those of Case 1 and Case 2. Each loading was represented by its peak angular acceleration magnitude in Figure 9. It can be noted that, for both concussive impacts, halving the acceleration magnitudes decreased 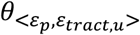 and the percentage values for the difference between 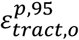 and 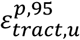. Conversely, both *θ_<ε_p_,ε_tract,u_>_* and the percentage values characterizing the variation between 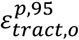 and 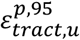 were elevated when doubling the acceleration profiles of two concussive impacts. Thus, the level of fiber orientation variation and its influence on the tract-oriented strain appear to be dependent on impact severity.

**Figure 9.**
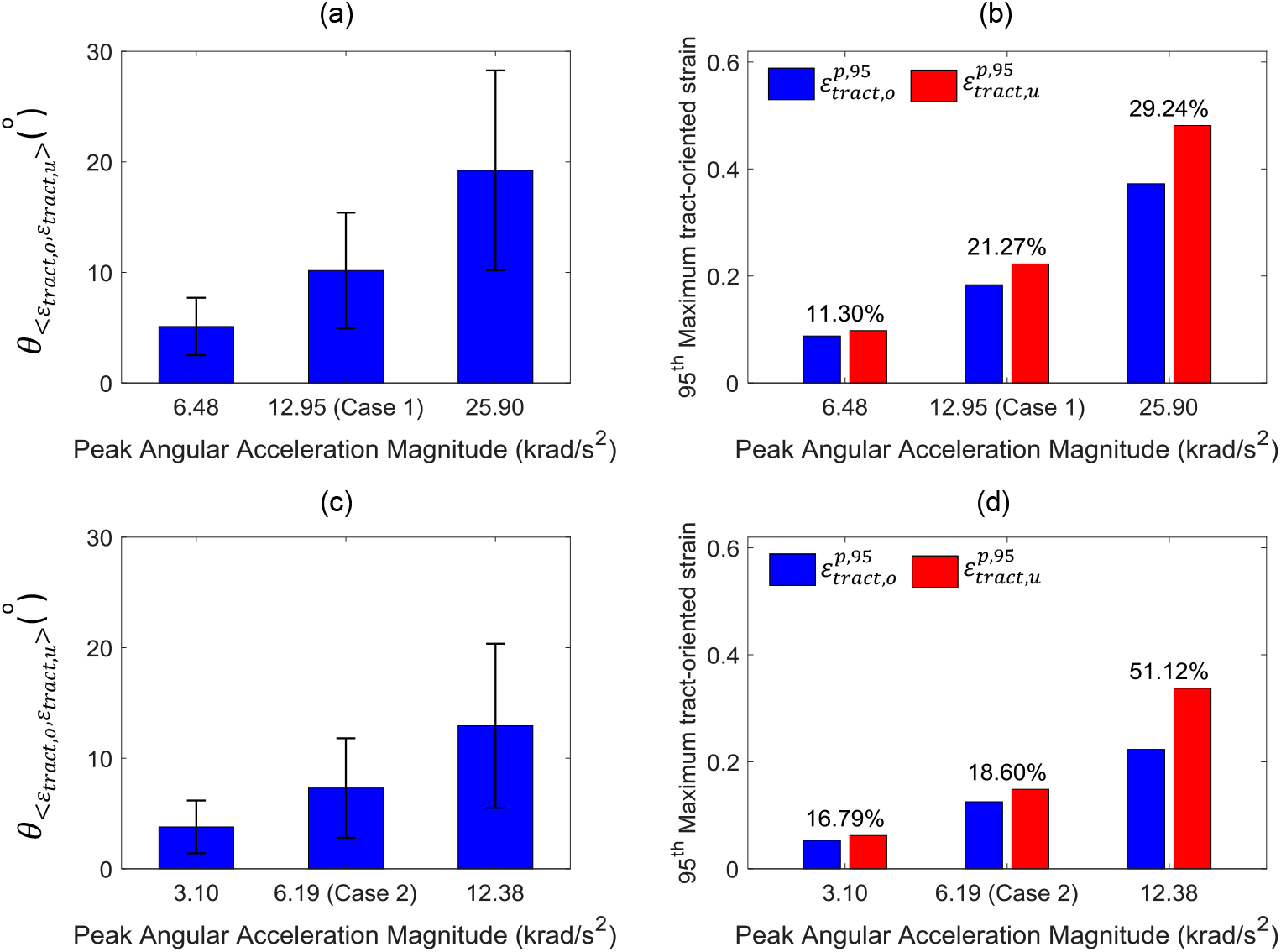
Influence of impact severity on fiber orientation variation and 95^th^ percentile maximum tract-oriented strain. (a) Element-wise *θ_<ε_tract,o_,ε_tract,u_>_* peak in the form of mean value ± standard deviation for Case 1 and two extra simulations with their loadings obtained by halving and doubling the acceleration profiles of Case 1, respectively, with the secondary effects on 95^th^ percentile maximum tract-oriented strain plotted in subfigure (b). (c) Element-wise *θ_<ε_tract,o_,ε_tract,u_>_* peak in the form of mean value ± standard deviation for Case 2 and two extra simulations with their loadings obtained by halving and doubling the acceleration profiles of Case 2, respectively, with the secondary effects on 95^th^ percentile maximum tract-oriented strain plotted in subfigure (d). Percentage values for the difference between 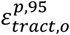 and 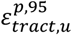 are listed in the bar figures. The impact severity was characterized by peak angular acceleration magnitude.

## Discussion

The current study detailed the implementation of an embedded element approach to monitor the real-time fiber orientation in a 3D head model, revealing the fiber orientation as a temporal variable during the impacts. A new scheme was proposed to calculate the tract-oriented strain by resolving the strain tensors from pre-computed simulations along the temporal fiber orientation instead of its static counterpart directly obtained from the DTI. It was verified that incorporating the real-time fiber orientation significantly increases the tract-oriented strain and consequently contributes a generally more extended distribution and a larger volume ratio of WM with high deformation along the fiber tracts.

Being an orientation-dependent parameter, the tract-oriented strain is highly sensitive to the variation of fiber orientation. By examining the relative angles between *ε_p_* and two types of tract-oriented strains, it was found that the fiber tracts tended to be reoriented toward the direction of *ε_p_* during the impacts. This tendency of directional alignment was also noted in an *in vitro* experimental model: by visualizing the configurations of nerve fibers within the brain tissue samples before and after applying uniaxial stretch, Tamura and colleagues^69^ observed that the directions of nerve fibers were typically oriented to be more parallel to the stretching direction. The congruency between the current study and the *in vitro* model increases the credibility of the presented computational results.

Besides quantifying the local deviation of fiber orientation, the current study also found that incorporating real-time fiber orientation amplifies the magnitude of the tract-oriented strain and the volume ratio of WM enduring high deformation along its axonal fiber orientation. This finding can be explained by the theory of continuum mechanics.^70^ A given strain tensor can be projected to arbitrary directions, resulting in different measures. The maximum principal strain (*ε_p_*) is an invariant of the strain tensor calculated as its maximum eigenvalue, characterizing the largest stretch of an elemental volume along the corresponding eigenvector (i.e., the direction of *ε_p_*). Thus, the more the fiber direction aligns with the eigenvector, the closer the magnitude of tract-oriented strain is to that of *ε_p_*. The tendency of directional alignment between the real-time fiber orientation and *ε_p_* provided a theoretical explanation for the elevated value observed from *ε_tract,u_* with respect to *ε_tract,0_*. Such results also suggested that the tract-oriented deformation may have bene underestimated in previous studies ^38, 41, 51^ in which the strain tensor was projected onto a time-invariant vector directly obtained from the DTI (similar to the scheme used to compute *ε_tract,o_* in the current study).

Possibly because of incorporating fiber direction, the tract-oriented strain has repeatedly been shown to be an appropriate mechanical parameter for TAI prediction.^37, 39, 41, 45, 48, 49^ Nevertheless, the majority of computational studies exclusively employed the accumulated maximum principal strain 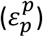 and its derivatives (e.g., CSDM, rate of *ε_p_*, the product of strain and strain rate) as injury predictors in TBI analysis. The current study proposed a new scheme to calculate the tract-oriented strain by incorporating the real-time fiber orientation (i.e., 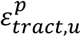). Thus, it is worth examining the relationship between the prevalently used 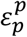 and the newly calculated 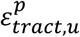. As illustrated in Figure 10, the model predicted larger 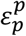 than 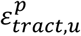 for both concussive impacts, consistent with previous findings^35, 43^ that the principal strain was an over-prediction of 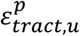 for brain tissue. By simultaneously scrutinizing the data in Figure 8 and Figure 10, no consistent scaling from 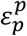 to 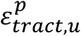 can be detected, indicating that the local strain in the axonal fiber cannot be trivially correlated to the maximum principal strain without taking into account the real-time fiber orientation. This further supports the necessity of incorporating the 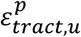 as the injury predictor in TBI analysis in the future, especially for TAI analysis.

**Figure 10.**
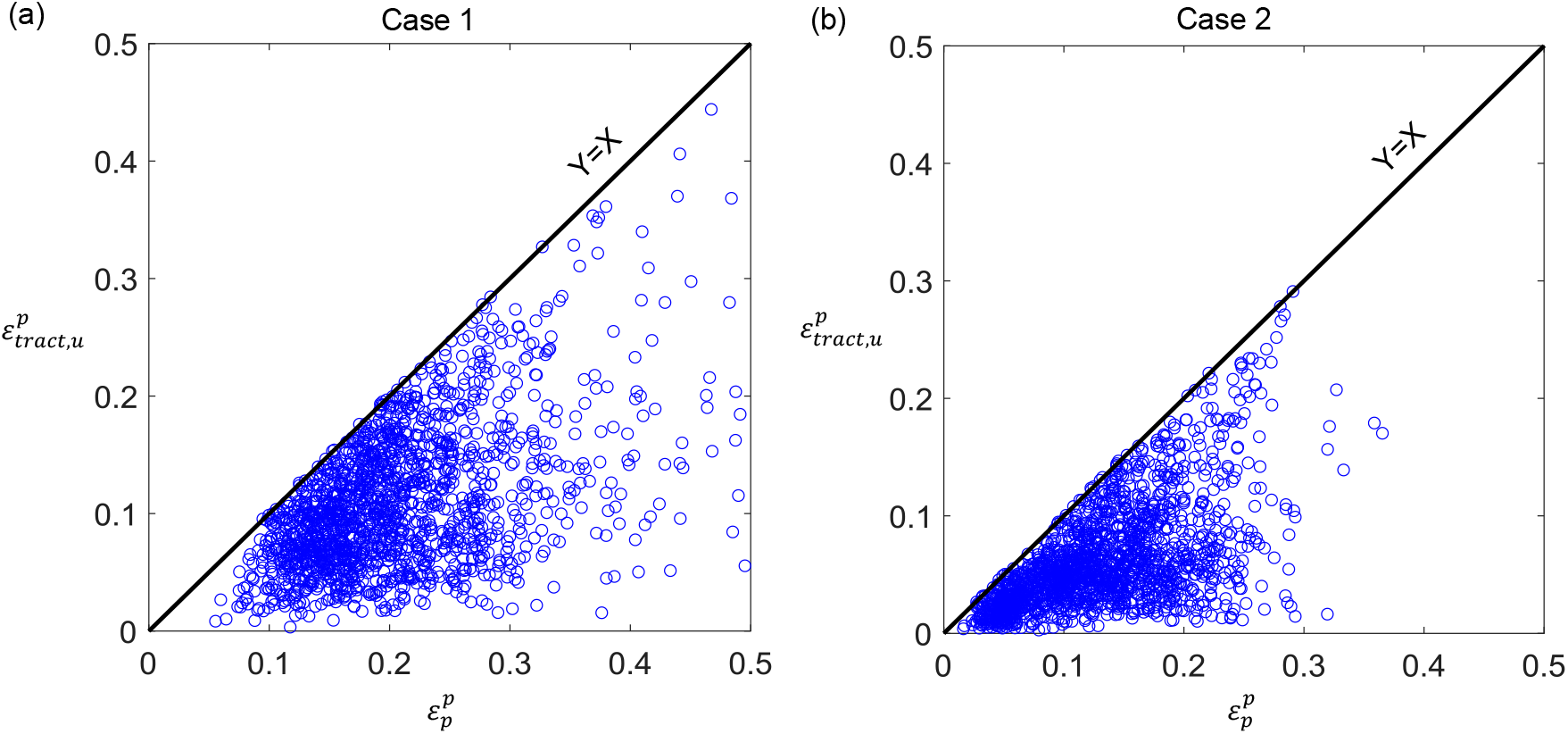
Element-wise relationship between 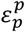 and 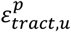 in Case 1 (a) and Case 2 (b).

The current study leveraged the embedded element approach to track the temporal fiber orientation during the simulations. As previously mentioned, the embedded truss elements only served as auxiliary for obtaining the updated orientation of fiber tracts without additional contribution to mechanical stiffness. Such a modeling strategy was deliberately designed to circumvent volume redundancy. In the embedded element approach, the master elements occupy the full volume of the model, including the volume of the embedded elements. Such an unphysical redundancy triggers a potential underestimation of strains and requires extra elimination.^35, 43^ In the current study, the truss elements were simulated as a null constitutive model with nominal density and cross-sectional area. Per the benefit of this modeling strategy, brain strain response was independent of the addition of truss elements given no differences were noted in the strain responses between the Baseline-model and the Truss-embedded-model (Figure 11). This finding further verified that volume redundancy was indeed avoided in the current study. It should also be clarified that the infinitesimal values for the material properties of truss elements (Table 1) were artificially defined and did not represent the structural dimension of the axonal fibers, similar to the strategy adopted by Wu and colleagues.^43^ This was valid given the truss element was intended to only track the real-time fiber orientation without additional mechanical contribution. Furthermore, the real-time fiber orientation was also considered in those studies that explicitly modelled the whole brain tractography,^35, 37, 43, 48^ but did not quantify the influence of real-time fiber orientation.

**Figure 11.**
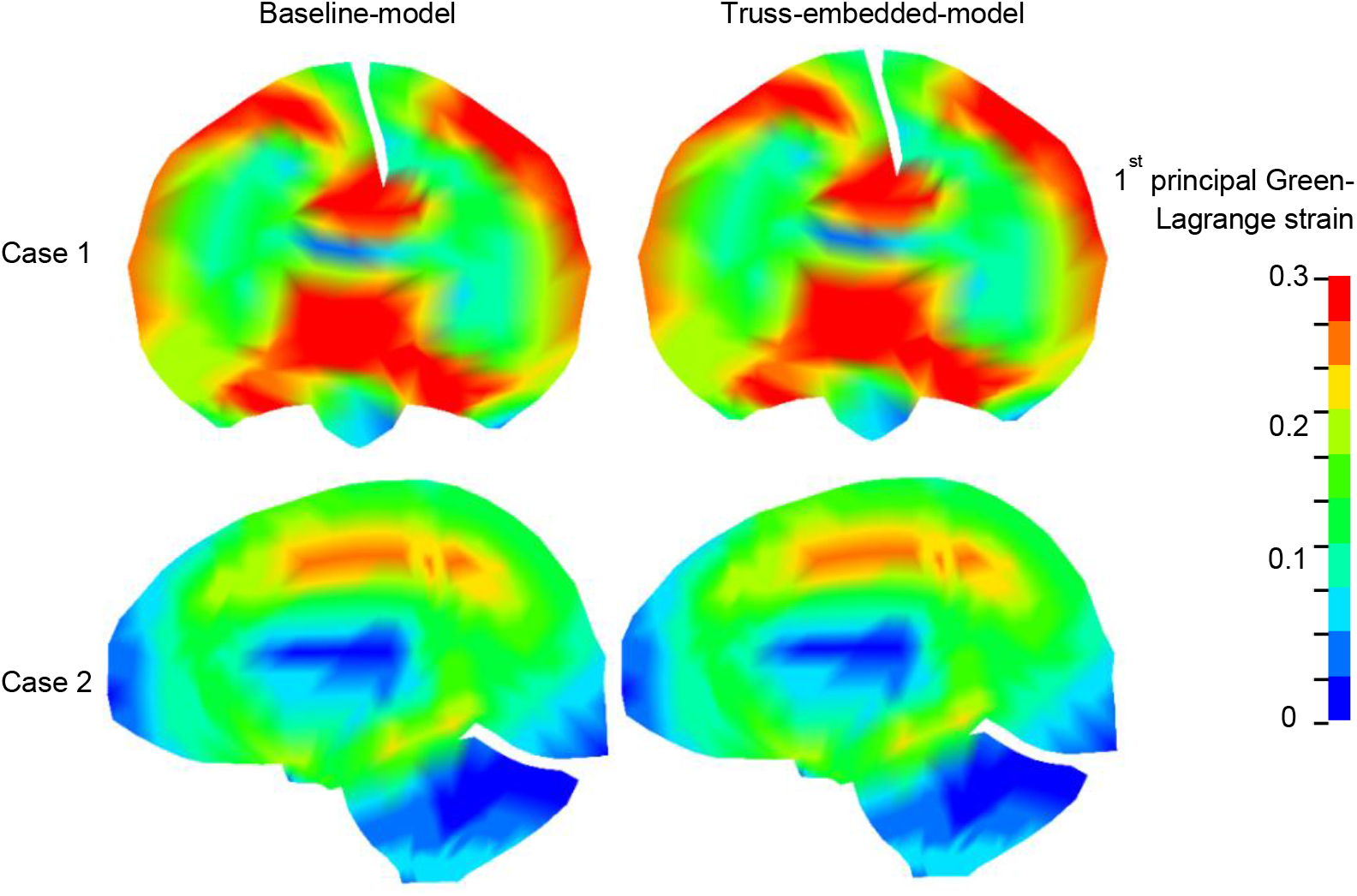
Comparison of *ε_p_* distribution in the brain between the Baseline-model and the Truss-embedded-model. Top row: Coronal cut-sections of *ε_p_* contour at 43 ms for Case 1. Bottom row: Sagittal cut-sections of *ε_p_* contour at 23 ms for Case 2.

The current study employed an isotropic constitutive law to represent the mechanical behavior of the brain tissue, similar to other works evaluating the tract-oriented strain.^39–41, 47, 71^ This modeling choice was supported by the findings that no significant mechanical dependency on fiber orientation was noted in the brain tissue.^72–75^ Nevertheless, the authors acknowledged that there were a large number of mechanical testing studies reporting that brain tissue was mechanically anisotropic.^76–80^ To date, several fiber-reinforced hyperelastic anisotropic constitutive models were implemented in the FE models to emulate the direction-dependence of the brain material, such as the Gasser-Ogden-Holzapfel model.^31, 32, 34, 36, 37, 42–44, 81^ As was in common in these studies, a correlation between fractional anisotropy (FA) from DTI and fiber-contributed stiffness was assumed. However, the validity of such modeling assumptions required further verification.^54, 82^ For those studies adopting the embedded element approach to explicitly represent the WM fiber tractography, the brain anisotropy can be alternatively simulated by assigning certain mechanical stiffness to the embedded elements, as done in the studies by Garimella and coworkers^35, 83^ and Wu and colleagues.^43^ As discussed above, this may raise the issue of volume redundancy, as is the case in the study by Garimella and Kraft.^35^ Lately, both Garimella and coworkers^83^ and Wu and colleagues^43^ appropriately addressed the volume redundancy by subtracting the constitutive contribution of the embedded elements from their counterparts of the master elements. Whereas, given the paucity of material properties and geometric features that were directly measured from the axonal fibers themselves, a direct calibration on the constitutive material model for the fiber tractography remains to be performed.^43^ Taken together, the authors acknowledged that no consensus has been reached yet, neither for the constitutive modeling of brain tissue nor for the mechanical contribution of axonal fibers in the FE models.^54, 84^

### Limitations and future work

Although this study yielded new insights about integrating the real-time fiber orientation into the calculation of the tract-oriented strain, certain limitations existed that requires further investigation. First and foremost, to ensure computational efficiency, an FE model with relatively low mesh density was used in the current study. To map the orientation information from the DTI with a resolution of 1 mm to the FE model with a mean mesh size of 5.2 mm, a weighted-average approach was used, which partially caused loss of orientation information and might have consequently affected the accuracy of the tract-oriented strain.^35, 71^ As detailed in the discussion, the strain-induced directional alignment between the real-time fiber orientation and that of *ε_p_* provided a theoretical explanation for the elevated value of *ε_tractu_* with respect to *ε_tract,o_*. The general agreement between the current study and a previous *in vitro* model^69^ further supported our advocation to incorporate the real-time fiber orientation in the calculation of the tract-orientation strain. In addition, the major finding revealed by the relatively coarse model in the current study was verified by another refined FE head model with a mean mesh size of around 1 mm,^40^ comparable to the DTI resolution. Detailed results from the refined FE head model will be presented in a future study focusing on the dependency of the tract-oriented strain on mesh size.

Compared to the studies^35, 37, 43^ in which the whole brain tractography was explicitly incorporated with continuous fiber tracts, the fiber orientation in the current study was downsampled given that only one fiber orientation was assigned to each WM element. Whereas, tractography was not considered to be a gold-standard of white matter architecture (in fact, there is no gold standard yet).^85^ Some of the known limitations of tractography include challenges in regions of crossing fibers with multiple orientations present within each voxel, which constitute a significant portion of WM. A future version of the proposed model^40^ with a mesh resolution that is comparable to that of DTI can increase the number of embedded truss elements, and potentially model fiber crossings by embedding multiple truss elements within one WM element.

Another potential limitation was that no relative motion between the truss elements and their master elements was permitted, similarly to the strategies adopted in other computational studies.^35, 37, 43^ In contrast to the current study, a gradual coupling between the undulated axons and the surrounding glial cells secondary to mechanical stretching was previously noted in the guinea pig optic nerves^86^ and the developing chick embryo spinal cord.^87^ Considering the inter-species variation, extrapolation of the results derived from animal experiments to human subjects needs further verification.^88, 89^ Moreover, the axonal architectures in the central nervous system exhibit region-dependency. For example, axonal undulation is less prominent in the cerebral region^90^ with respect to the optic nerves and spinal cord. Such a region-dependency further warrants caution regarding the extrapolation of results obtained from the optic nerves and spinal cord to the cerebrum, cerebellum and brainstem as the regions of interest in the current study.

The current study examined the dependency of fiber orientation variation and tract-oriented strain on impact severity with only three levels of acceleration magnitude incorporated. A systematic investigation that covers more impact-related variables (e.g., rotational velocity, impact duration, rotational directions) with their magnitudes spanning over the regimes measured from the realistic impacts is planned for future work to identify the critical scenarios that the fiber orientation exhibited a more pronounced effect. In addition, it should be further stressed that the current study was conducted based on the assumption of the brain as an isotropic medium. Extrapolating of findings in the current study to those that regarded the brain as an anisotropic structure requires further investigation, which could be another possible direction for future work.

## Conclusion

The present study employed an embedded element approach to monitor the real-time fiber orientation during head impacts and proposed a new scheme to calculate the tract-oriented strain by projecting the strain tensors from pre-computed simulation along the temporal fiber direction instead of its static counterpart directly from the DTI. It was revealed that incorporating the real-time fiber orientation not only altered the direction but also amplified the magnitude of the tract-oriented strain, leading to a generally more extended distribution and a larger volume ratio of WM exposed to high deformation along the fiber tracts. These effects were exacerbated with the impact severities as characterized by the acceleration magnitude. This study provides guidance on how best to incorporate the fiber orientation in head injury models and derive the tract-oriented strain from computational simulations.

## Acknowledgements

Drs. Michael Zeineh, Gerald Grant, and David Camarillo received funding from the Pac-12 Conference’s Student-Athlete Health and Well-Being Initiative and Taube Stanford Children’s Concussion Initiative. The content of this article is solely the responsibility of the authors and does not necessarily represent the official views of funding agencies. The authors would like to thank Dr. Chiara Giordano for providing an earlier version of the Python code, which was adapted and further used for the computation of the tract-oriented strain in this study. The technical support provided by Dr. Hao Chen from Livermore Software Technology Corporation, Livermore, CA (US) is gratefully acknowledged. The authors thank the Swedish Research Council (VR-2016-05314) for the support. The simulations were performed on resources provided by the Swedish National Infrastructure for Computing (SNIC) at the center for High Performance Computing (PDC). The authors also thank Dr. Jean Ann Sanford and the anonymous reviewer for the stimulating comments and valuable suggestion that substantially improved this paper. The help from Ms Nicole Mouchawar for proofreading is highly appreciated.

## Appendix A: Sensitivity analysis on the length of embedded truss elements

A sensitivity analysis was conducted to investigate the dependency of fiber orientation variation (i.e., *θ_<ε_tract,o_,ε_tract,u_>_*) and strain response on the length of the embedded truss element. For the strain response, the first principal Green-Lagrange strain (i.e., *ε_p_*) and the tract-oriented Green-Lagrange strain (characterized by 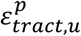 and 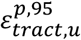) were selected, both of which were parameters of interest in the current study. A detailed definition for these symbols is available in Table 3 in the “Post-processing and data analysis” section.

To achieve this analysis, three models were developed with the length of embedded truss elements as 2 mm in Model 1, 1 mm in Model 2, and 0.5 mm in Model 3 (Figure A1). For a given truss element that was correspondingly embedded within a white matter (WM) element (serving as a master element), one node located at the centroid of the master element and the other fell within the boundary of the master element. The orientation of the truss element was obtained from the diffusion tensor imaging (DTI) via a weighted-average procedure as detailed in the “Computation of the tract-oriented strain with static fiber orientation” section. Material properties of the truss elements and the coupling strategy between truss elements and WM elements were the same as those detailed in the “Implementation of embedded element method for fiber orientation modeling” section. Thus, the only difference among the three models was the length of the truss elements. A concussive impact (i.e., Case 1 as detailed in the “Loading conditions” section) was imposed on three models, respectively.

**Figure A1.**
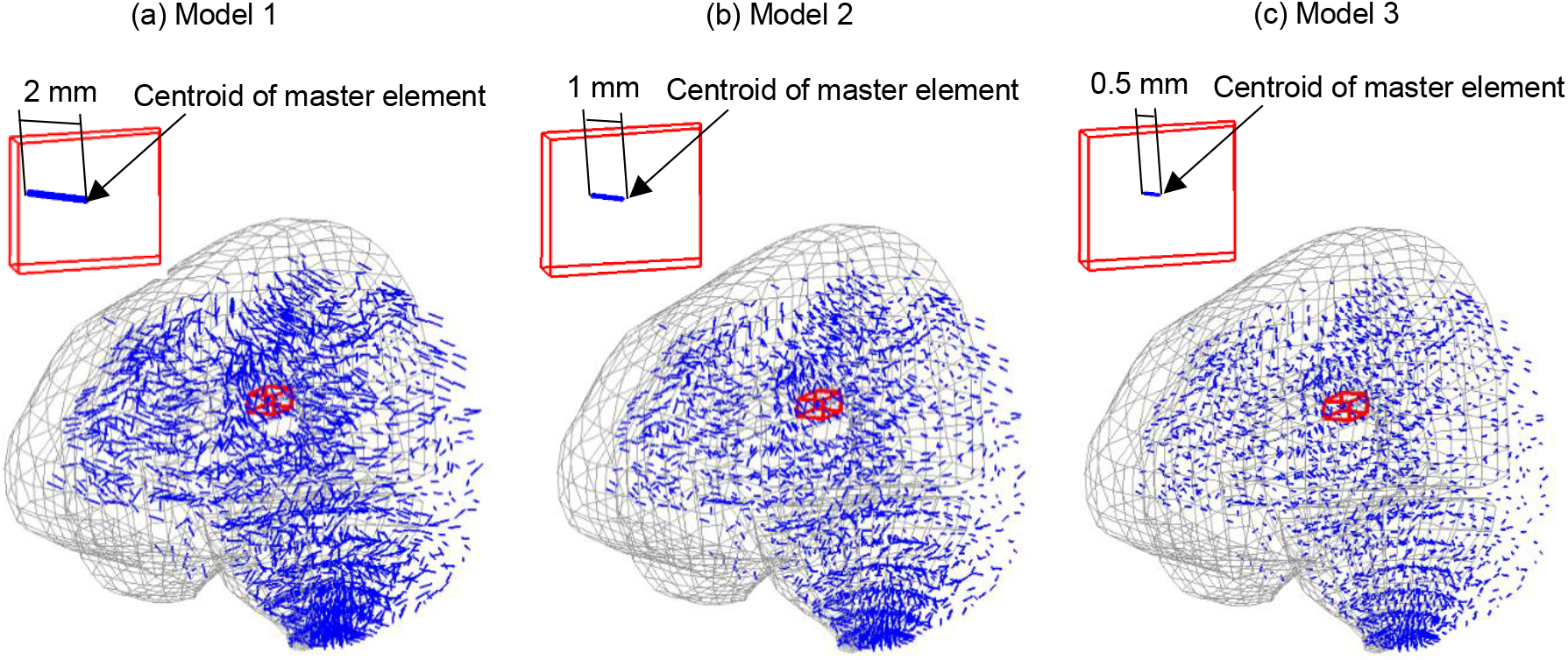
(a) Isometric view of the brain (in gray) in Model 1 with the embedded truss elements as 2 mm (in blue). (b) Isometric view of the brain (in gray) in Model 2 with the embedded truss elements as 1 mm (in blue). (c) Isometric view of the brain (in gray) in Model 3 with the embedded truss elements as 0.5 mm (in blue). In all subfigures, half of the brain is masked for better illustration and an enlarged view is provided for a representative element to illustrate the length of the truss element.

Figure A2 plots the time-history curves of maximum *θ_<ε_tract,o_,ε_tract,u_>_* in six brain subregions. It can be noted that the maximum *θ_<ε_tract,o_,ε_tract,u_>_* was insensitive to the length of the truss elements, given that the curves predicted by the three models overlapped.

**Figure A2.**
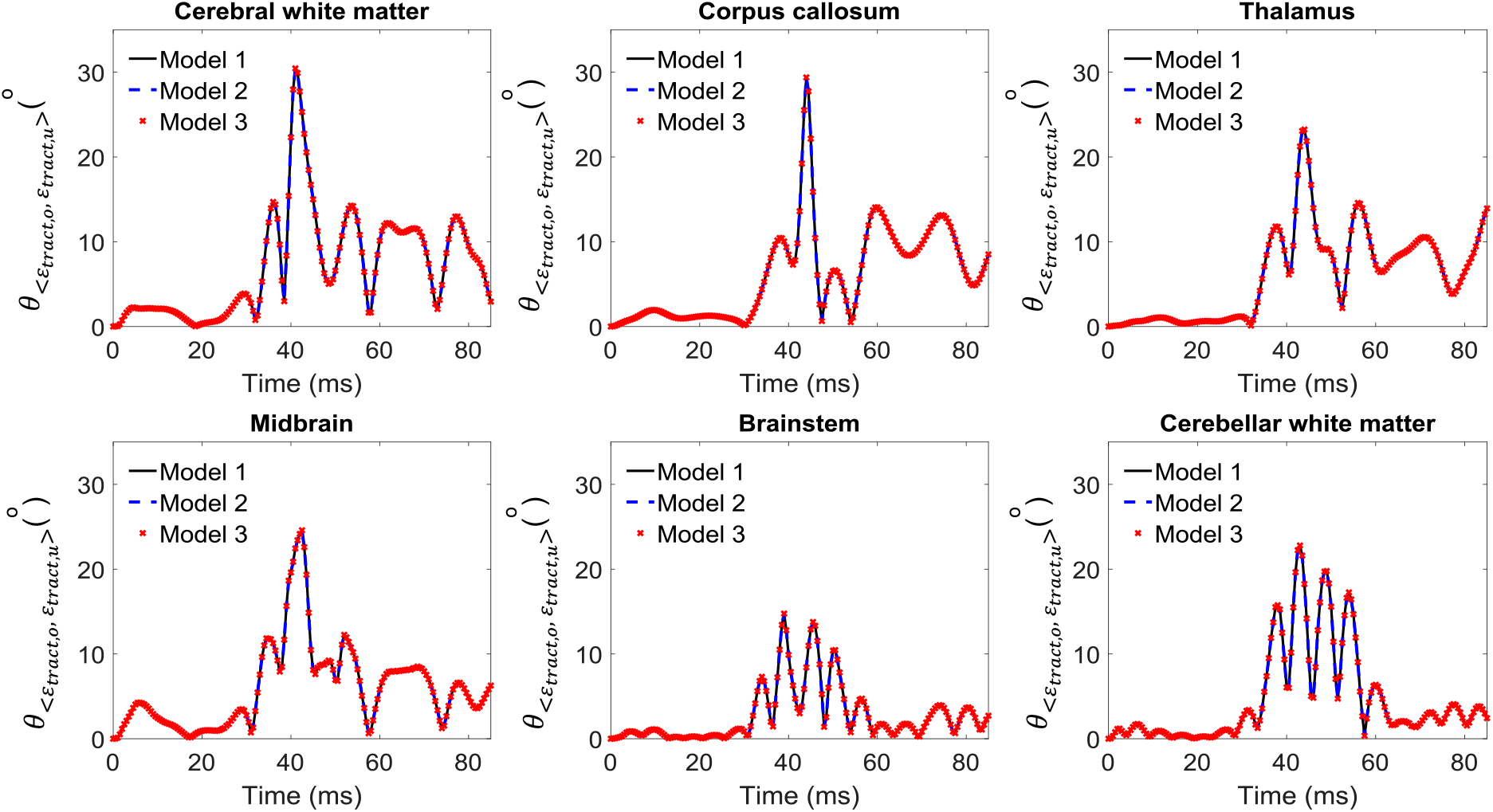
Time history curves of maximum *θ_<ε_tract,o_,ε_tract,u_>_* in six brain subregion predicted by the three models for Case 1.

Figure A3 shows the coronal cut-section *ε_p_* contour without notable variation among the three models. For the tract-oriented strain, the three models predicted the identical value of 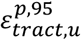 as 0.22 (Figure A4 (a)). Same distributions of WM elements with 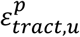 over 0.14 were also noted among the three models (Figure A4 (b)).

**Figure A3.**
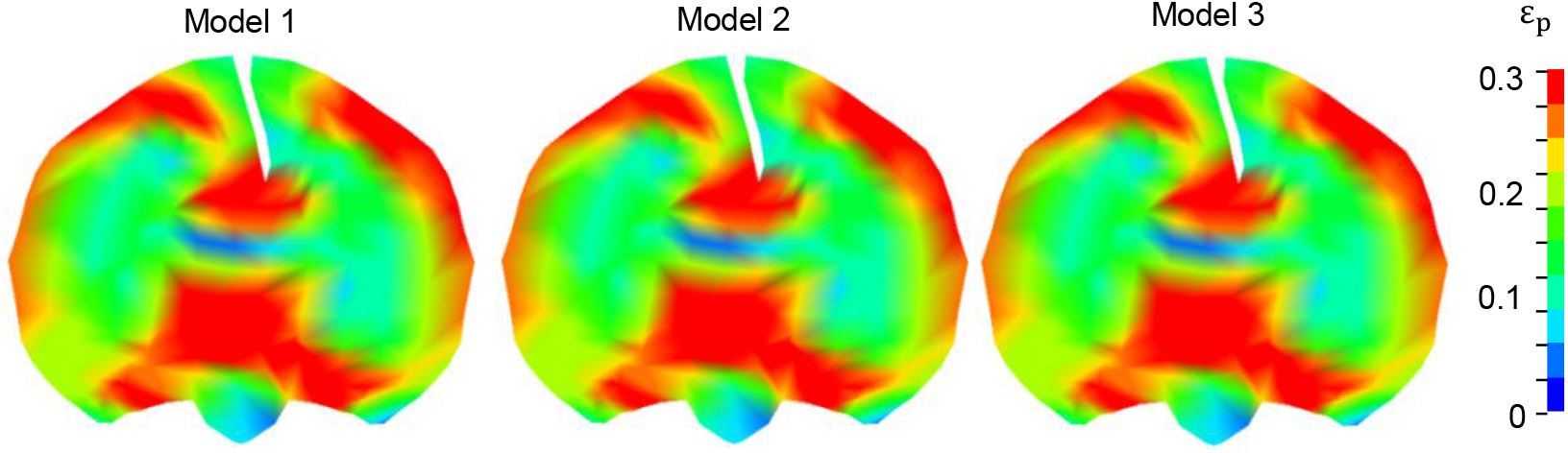
Coronal cut-section of *ε_p_* contour predicted by the three models at 43 ms for Case 1.

**Figure A4.**
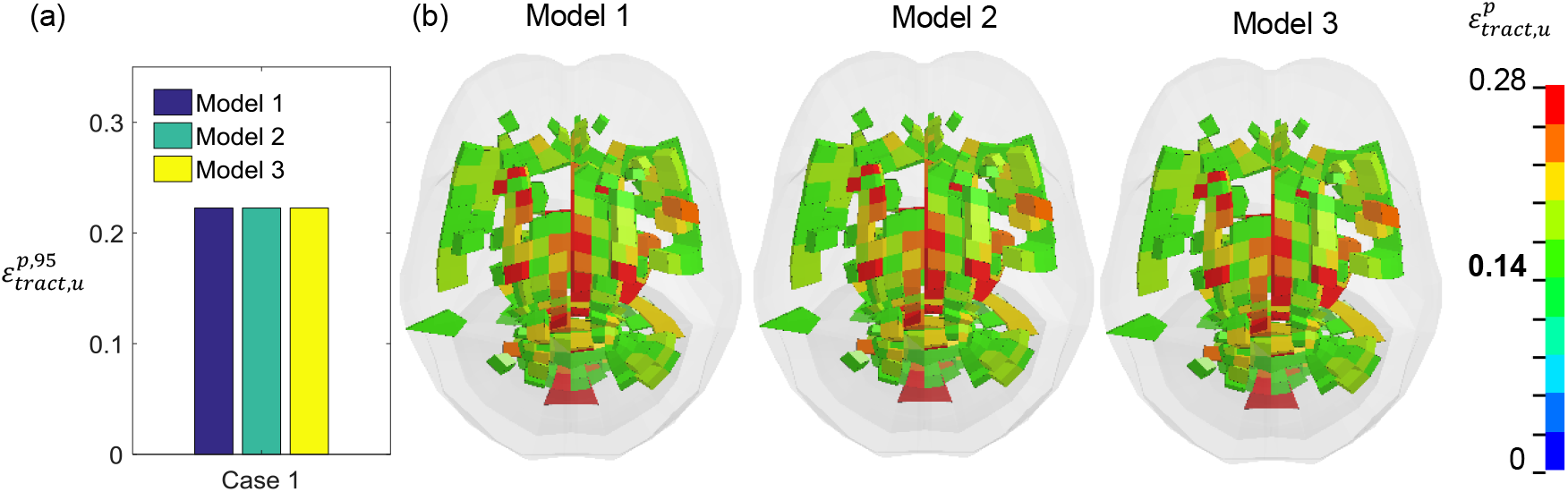
(a) Comparison of 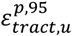 predicted by the three models for Case 1; (b) Top view of WM elements with 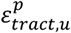 over 0.14 for Case 1.

Based on the identical response from the three models, it can be concluded that, under the condition that a truss element oriented along a predefined direction with both ends within the same master element, neither the deviation of fiber orientation nor the brain strain response was sensitive to the length of the embedded truss elements.

